# Cuticular profiling of insecticide resistant *Aedes aegypti*

**DOI:** 10.1101/2023.01.13.523989

**Authors:** Ella Jacobs, Christine Chrissian, Stephanie Rankin-Turner, Maggie Wear, Emma Camacho, Jeff G. Scott, Nichole A. Broderick, Conor J. McMeniman, Ruth E. Stark, Arturo Casadevall

## Abstract

Insecticides have made great strides in reducing the global burden of vector-borne disease. Nonetheless, serious public health concerns remain because insecticide-resistant vector populations continue to spread globally. To circumvent insecticide resistance, it is essential to understand all contributing mechanisms. Contact-based insecticides are absorbed through the insect cuticle, which is comprised mainly of chitin polysaccharides, cuticular proteins, hydrocarbons, and phenolic biopolymers sclerotin and melanin. Cuticle interface alterations can slow or prevent insecticide penetration in a phenomenon referred to as cuticular resistance. Cuticular resistance characterization of the yellow fever mosquito, *Aedes aegypti*, is lacking. In the current study, we utilized solid-state Nuclear Magnetic Resonance (ssNMR) spectroscopy, gas chromatography/mass spectrometry (GC-MS), and transmission electron microscopy (TEM) to gain insights into the cuticle composition of congenic cytochrome P450 monooxygenase insecticide resistant and susceptible *Ae. aegypti*. No differences in cuticular hydrocarbon content or phenolic biopolymer deposition were found. In contrast, we observed cuticle thickness of insecticide resistant *Ae. aegypti* increased over time and exhibited higher polysaccharide abundance. Moreover, we found these local cuticular changes correlated with global metabolic differences in the whole mosquito, suggesting the existence of novel cuticular resistance mechanisms in this major disease vector.

## Introduction

Worldwide, vector control programs rely on insecticides to prevent vector-borne diseases. Mosquitoes are responsible for the most significant burden of vector-borne disease and consequently are a primary focus of public health interventions. *Aedes aegypti* is the primary vector of four arboviruses: yellow fever (YFV), dengue (DENV), chikungunya (CHIKV), and Zika (ZIKV) [1]. Of these arboviruses, dengue virus has the most significant public health burden with an estimated 50-100 million symptomatic infections per year [2, 3]. Moreover, *Ae. aegypti* has an encroaching geographic range that renders half of the world’s population at risk for dengue infection [4]. Vector control in dengue endemic areas relies heavily on pyrethroid insecticides [5-8]. Over time, the selection pressure imposed by the extensive use of pyrethroids in these regions has resulted in increasingly widespread populations of highly insecticide resistant *Ae. aegypti* [7]. Consequently, insecticide resistance threatens to undermine *Ae. aegypti* control efforts in regions where vector-borne diseases are most prevalent.

Insecticide resistance is complex and often involves physiological resistance and behavioral responses [9-12]. Physiological resistance occurs via two primary mechanisms: target-site mutations and metabolic resistance. Contact-dependent pyrethroid insecticides target the *voltage-sensitive sodium channels* (*Vssc*), which are essential to the insect nervous system [13]. Insects with mutations in *Vssc* that prevent pyrethroid activity are phenotypically *knockdown resistant* (*kdr*) [14, 15]. Metabolically resistant insects exhibit increased expression and detoxifying activity of cytochrome P450 monooxygenases (CYPs) allowing for metabolization and excretion of insecticides [16, 17]. A large body of work has demonstrated that these physiological adaptations play a significant role in insecticide resistance [11, 16]. In contrast, however, comparatively little is known about other physiological mechanisms that contribute to resistance, such as modifications to the insect exoskeleton [11, 16, 18]. The exoskeleton, also referred to as insect cuticle, is essential for structural integrity, barrier protection, sensation, hydration, and chemical communication [19, 20]. Furthermore, the cuticle is an arthropod’s first line of defense against contact insecticides [18]. Modifications to the cuticle including thickening have been shown to slow, or even prevent the penetration of contact insecticides — a phenomenon first documented in the 1960s — yet the structural modifications and mechanisms contributing to cuticular thickening remain largely uncharacterized [11, 18, 21, 22]. Although cuticular resistance alone is insufficient to confer complete resistance, it acts synergistically with other resistance mechanisms to promote efficient insecticide elimination and limit internal damage [16, 18].

Mosquito cuticle is a formidable barrier to external assault; it is composed of three distinct layers, each with their own unique properties. The epicuticle is a thin, waxy, hydrocarbon-rich layer deposited on the outermost surface of the cuticle. Beneath the epicuticle lies the exocuticle, followed by the endocuticle. Both the exo- and endocuticular layers consist of macromolecular frameworks composed of the polysaccharide chitin, with proteins and lipids interwoven throughout [20, 23]. The exocuticle has a distinct lamellar structure that forms soon after adults eclose from pupae, whereas the soft endocuticle is deposited after eclosion [23, 24]. The lamellar structure of the exocuticle is hard and rigid due to the deposition of tyrosine-derived sclerotin, a process known as sclerotization. During sclerotization, chitin is cross-linked to key residues of cuticular proteins via the oxidative conjugation of tyrosine-derived catechols, resulting in cuticular hardening conferring structural stability and resiliency. The pigmented biopolymer melanin, another tyrosine-derived component of the exocuticle, is responsible for cuticular darkening [25-29]. Like sclerotin, melanin is a highly recalcitrant phenolic-based polymer that is crosslinked to other cuticular moieties, albeit to a lesser degree [23, 30, 31]. Melanin has several outsized roles in other physiological processes such as wound healing and the insect immune response [28]. Phenoloxidases, including laccases and tyrosinases, produce phenolic biopolymers for these various processes under tight regulation to limit internal damage [32, 33]. Due to their compatibility with a range of substrates, phenoloxidases have a proposed role in detoxification in many invertebrates [34-36]. While sclerotin and melanin are present in insect cuticle in relatively small quantities, they are biologically important and their presence in the cuticle suggests that these two phenolic biopolymers may contribute to cuticular insecticide resistance.

Recent work has demonstrated that cuticles of insecticide resistant populations in malaria vectors *Anopheles gambiae* and *An. funestus* possess distinct structural and biochemical alterations [18]. In contrast, very few studies have directly characterized cuticular alterations in the major arboviral vectors of the genus *Aedes* [37, 38]. One notable feature of this resistant phenotype is leg cuticular thickening. Mosquitoes are often exposed to insecticides while resting on treated surfaces, rendering the leg cuticle an important interface for contact-dependent insecticide absorption [12, 39, 40]. In comparisons of mosquito leg cross-sections using electron microscopy (EM), insecticide resistant *An. gambiae* and *An. funestus* were found to have thicker cuticles [40-44]. While EM is well-suited to characterize the exo- and endocuticular layers, the thin, waxy hydrocarbon-rich epicuticle is not commonly visible [40]. However, the long-chain (∼C21–C37+) alkane or alkene cuticular constituents are readily extractable from the epicuticle and can be profiled using gas chromatography/mass spectrometry (GC-MS) [45]. These aliphatic lipids have been shown to serve as a barrier to desiccation, and their increased cuticular abundance has been correlated with insecticide resistance [19, 44, 46, 47].

No investigations of the potential contribution of sclerotin and melanin to cuticular resistance have appeared in the literature. More broadly, the phenomenon of cuticular resistance is understudied in *Ae. aegypti*. To address these shortcomings, we profiled cuticular differences between two characterized congenic strains: susceptible Rockefeller (ROCK) and CYP-mediated metabolically resistant strain herein referred to as CR [17]. This strain was derived from the well-characterized pyrethroid resistant Singapore (SP) strain that possesses resistance loci conferring both KDR and CYP resistance [16]. It is well-established that KDR and CYP interact with one another in nonadditive ways [48, 49]. To study the resistance mechanisms individually, the SP resistance loci were crossed into the ROCK genetic background and then isolated into two separate strains containing either the CYP-mediated resistance locus (CR) or KDR resistance locus [50]. The CR strain used in this work is primarily resistant to pyrethroid insecticides, with some cross-resistance to several organophosphate insecticides due to overexpression of several identified CYP genes [17, 50]. The possibility that the CR strain possessed cuticular morphology and compositional changes associated with the isolation of CYP-mediated resistance was unknown.

In this work we have utilized complementary biophysical, biochemical, and imaging methodologies to characterize the cuticle of the congenic ROCK and CR strains. Our studies focused primarily on female mosquitoes of this strain in light of the male’s inability to transmit vector-borne diseases. We adapted a methodology used to enrich for deposited fungal melanin to compare insecticide resistant and susceptible *Ae. aegypti* females [51]. Because insects produce sclerotin in addition to melanin, we utilized *Drosophila melanogaster* pigmentation mutants to validate the ability of this method to consistently recover phenolic compounds as well as their associated lipid and polysaccharides. Due to the structurally complex, insoluble nature of insect cuticle, we utilized solid-state Nuclear Magnetic Resonance (ssNMR) spectroscopy to compare the sclerotin and melanin-rich acid-resistant material as well as whole intact mosquitoes from insecticide resistant and susceptible *Ae. aegypti* females. Because the relationship between insecticide resistance mechanisms and phenoloxidase activity has not been fully elucidated [49, 52], we compared phenoloxidase activity in the two strains. Additionally, we monitored cuticular thickening and changes in cuticle ultrastructure over time using TEM at 3-5 d and 7-10 d post-eclosion. Finally, we analyzed the cuticular hydrocarbon content from both males and females using gas chromatography/mass spectrometry (GC-MS). Our analyses finds that neither cuticular hydrocarbons nor phenolic biopolymer deposition differed between insecticide resistant and susceptible mosquito strains. However, increased endocuticle deposition was observed in resistant mosquitoes, suggesting the existence of novel mechanisms of cuticular resistance in this globally important disease vector.

## Results

### Differences in phenoloxidase activity between CR and ROCK females

Phenoloxidases produce melanin pigments, which are important parts of the insect immune response. Constitutively active phenoloxidase activity, a measurement of baseline immune activation [53], was estimated by incubating individual female mosquito homogenates with 2 mM of the melanin precursor L-DOPA. There was no difference in constitutively active phenoloxidase between the two strains, indicating a similar level of baseline immune activation (Fig. 1A). However, because active phenoloxidases produce damaging reactive oxygen species, most phenoloxidases are present as zymogenic pro-phenoloxidases that require protease cleavage for activation [53, 54]. To estimate total phenoloxidase content, including zymogenic pro-phenoloxidases that require proteolytic cleavage, homogenates were incubated with the proteolytic enzyme α-chymotrypsin. A clearly diminished level of total phenoloxidase activity was evident in the insecticide resistant CR strain (Fig 1B).

**Figure 1:**
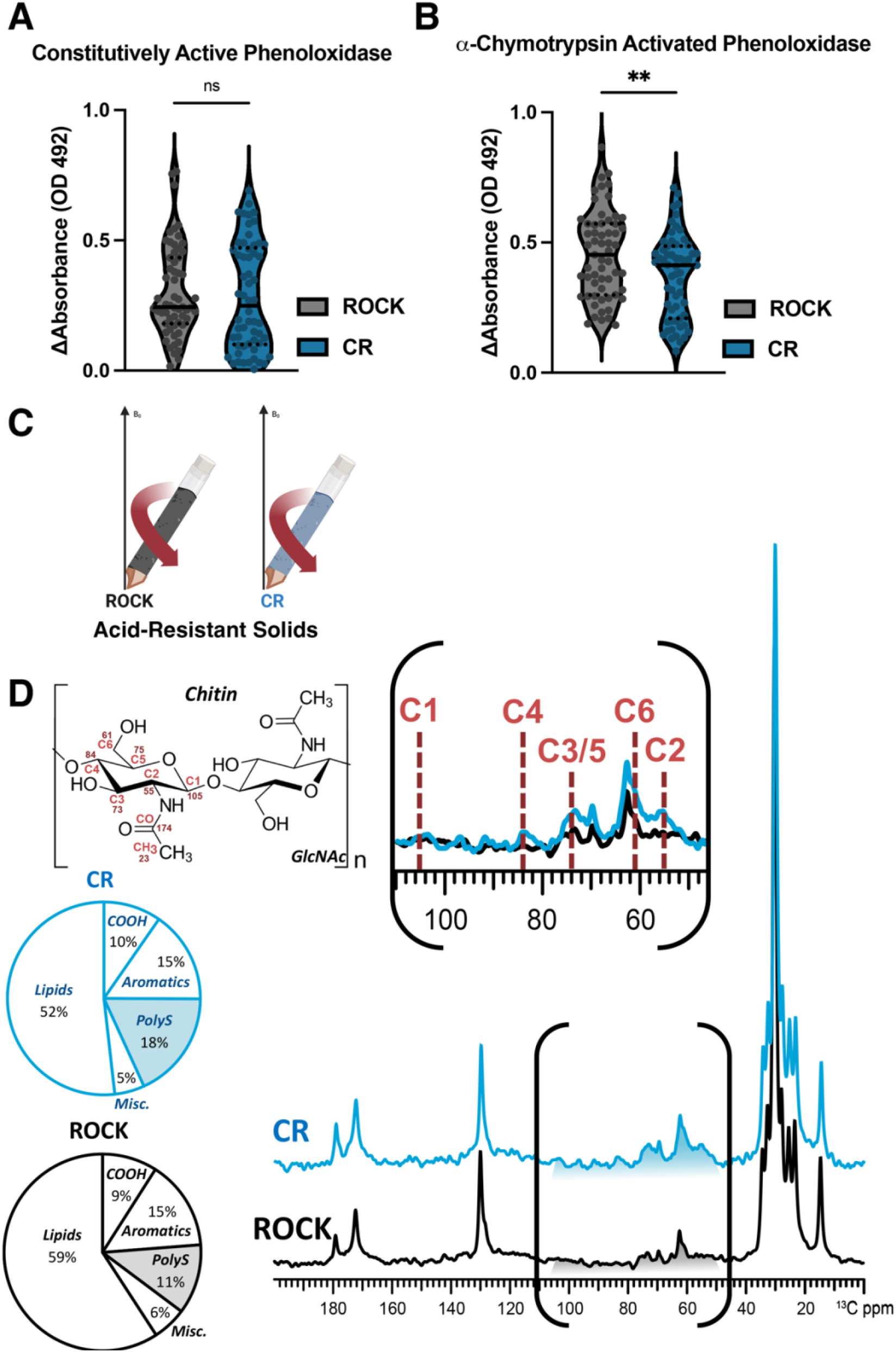
Characterization of phenoloxidase activity and cuticular content of insecticide resistant and susceptible *Ae. aegypti*. **A:** Constitutive phenoloxidase activity of insecticide-susceptible ROCK (grey) and insecticide-resistant CR (dark blue) female mosquito homogenate after incubation with 2 mM L-DOPA. Data points represent the change in absorbance (OD 492) after 45 minutes at 30°C from individual mosquitoes from three biological replicates. Sample size: ROCK n = 59 CR n = 58. Constitutively active phenoloxidase Mann-Whitney test p value = 0.3927 **B:** Constitutive phenoloxidase activity of insecticide-susceptible ROCK (grey) and insecticide-resistant CR (dark blue) female mosquito homogenate after incubation with 2 mM L-DOPA and 0.07 mg/mL *a*-chymotrypsin. Data points represent the change in absorbance (OD 492) after 45 minutes at 30°C from individual mosquitoes from three biological replicates. Sample size: ROCK n = 59 CR n = 58. *a*-chymotrypsin activated phenoloxidase unpaired students t-test p value = 0.0076. **C:** Schematic of solids loaded into ssNMR rotor to compare acid-resistant material from both strains **D:** ^13^C DPMAS ssNMR (50-sec delay; quantitatively reliable) comparison of acid-resistant material of the CR (dark blue) and ROCK (grey) strains pooled from three biological replicates.

### Validation of acid-resistant material from *D. melanogaster*

In addition to immunity, insect melanin, along with sclerotin, plays a vital role in maintaining the structural integrity of the cuticle [27]. Both melanin and sclerotin are found in the exocuticle, where they form strong covalent crosslinks to other cuticular moieties, such as chitin and proteins. We wanted to know whether these structurally amorphous phenolic biopolymers and their crosslinked constituents contribute to cuticular resistance. Prolonged HCl hydrolysis has been carried out previously on insects to isolate the acid-resistant portion of the cuticle, a method also implemented for the crude isolation of melanin deposited in fungal cells and mammalian hair [51, 55, 56]. Melanin and sclerotin share many biophysical properties, including acid degradation resistance, and therefore are virtually indistinguishable using these treatments [56, 57]. As such, it remained unclear whether the acid-resistant material yielded by the prolonged HCl digestion of insects was enriched in melanin, sclerotin, or a mixture of the two biopolymers. To verify that both melanin and sclerotin protect bonded constituents from acid hydrolysis, *D. melanogaster* flies from the pigmentation mutant strains ‘Ebony’ and ‘Yellow,’ which are unable to produce sclerotin and eumelanin, respectively, were subjected to prolonged digestion in concentrated HCl. The mass of the solid material recovered from each *D. melanogaster* strain, expressed as a percentage of the starting sample mass, is shown in Supplementary Fig. 1A. The eumelanin-deficient yellow strain and sclerotin-deficient ebony strain yielded acid-resistant material, but each of the two pigmentation mutant strains yielded less acid resistant material in comparison to the wild-type *D. melanogaster* strain (Yellow: 6.48%; Ebony: 7.80%; WT: 9.04%). These findings indicate that both sclerotin and melanin contribute to the acid resistance of insect cuticle. To confirm that prolonged HCl digestion was a suitable means to prepare insect cuticle samples that are enriched in both polymers, the acid-resistant material from each *D. melanogaster* strain was analyzed using Carbon-13 (^13^C) ssNMR spectra (Supplementary Fig. 1B). As anticipated, all three spectra were largely similar; they each displayed a broad resonance that spanned the aromatic carbon region (∼110-160 ppm) that is characteristic of amorphous phenolic polymers [58, 59] and contributed similarly to the overall signal intensity of each spectrum (Ebony and WT: 22.2%; Yellow: 22.1%).

### ssNMR analysis of acid-resistant material from CR and ROCK females

To our knowledge, no prior reports have analyzed phenolic biopolymers contribution to cuticular resistance. Although the masses of acid-resistant material were the same between the susceptible and resistant *Ae. aegypti* strains (Supp. Fig. 1A), we were curious to determine the chemical composition of these materials was unknown. To address this gap in knowledge, ^13^C ssNMR was performed to characterize the molecular architecture of the acid-resistant material recovered after HCl digestion of CR or ROCK mosquitoes (Fig. 1C). To probe for compositional differences between the two strains, direct-polarization (DPMAS) measurements with a long recycle delay (50 sec between successive data acquisitions) were carried, yielding data with quantitatively reliable peak intensities (Fig. 1D). Thus, the area of each spectral region compared to the total integrated area across the spectrum represents the relative amount of the corresponding acid-resistant cuticular moiety in the sample. The ^13^C ssNMR spectra of the acid-resistant samples from CR (blue trace) and ROCK (black trace) mosquitoes are shown in Figure 1D, each normalized to the tallest peak of the spectrum (∼30 ppm). The two spectra are generally similar in appearance: both display signals in the regions attributable to long aliphatic chains of hydrocarbons (∼10-40 and 130 ppm), polysaccharides (50-110 ppm) and pigments (110-160 ppm). The integrated area of the spectral region where the aromatic pigment carbons resonate contributes similarly to the total integrated signal intensity of each spectrum (15% for each), indicating that the relative amounts of phenolic biopolymers are similar in both samples.

However, by setting the tallest peak to full scale, a difference in the relative polysaccharide content between the two samples became apparent visually. This finding was corroborated by quantitative analysis: a relative increase in the polysaccharide content was observed in CR compared to ROCK (18% vs. 11%, respectively) with a concurrent decrease in lipid content (52% vs. 59%). Due to the acid digestion process and lack of phenolic biopolymers in the epicuticle, the lipids retained in this material are unlikely to be epicuticular hydrocarbons that can be extracted for GC-MS analysis. The polysaccharide content of these samples is likely to consist primarily of chitin, which is crosslinked to other cuticular moieties, because polysaccharides that are not covalently bonded within the cuticle are unlikely to withstand 24-hour digestion in concentrated HCl [60, 61]. This supposition is supported by chemical shift analysis of the ssNMR data. Whereas the ^13^C chemical shifts of the ring carbons of most polysaccharide species lie between 60 and 110 ppm, the C2 ring carbon of chitin, which is amide bonded to the acetyl-group nitrogen, gives rise to a characteristic ∼55 ppm signal that is clearly observed with greater intensity in the CR sample spectrum (Fig. 1D inset).

### Size and polysaccharide differences in insecticide resistant CR females

Previous work has demonstrated that CYP-mediated resistance in *Ae. aegypti* occurs with a trade-off in body size [49]. In agreement with previous findings, the CR female mosquitoes had significantly smaller body mass compared to ROCK females (Fig. 2A). However, it has not previously been assessed whether this difference in body mass is correlated with differences in the overall molecular composition of the whole insect, which could contribute to cuticular changes. Profiling of whole mosquitoes provides the opportunity to measure the polysaccharide content of the whole organism in addition to the lipids retained within insect fat bodies, which are also distinct from epicuticular hydrocarbons that can be extracted for GC-MS analysis [62].

**Figure 2:**
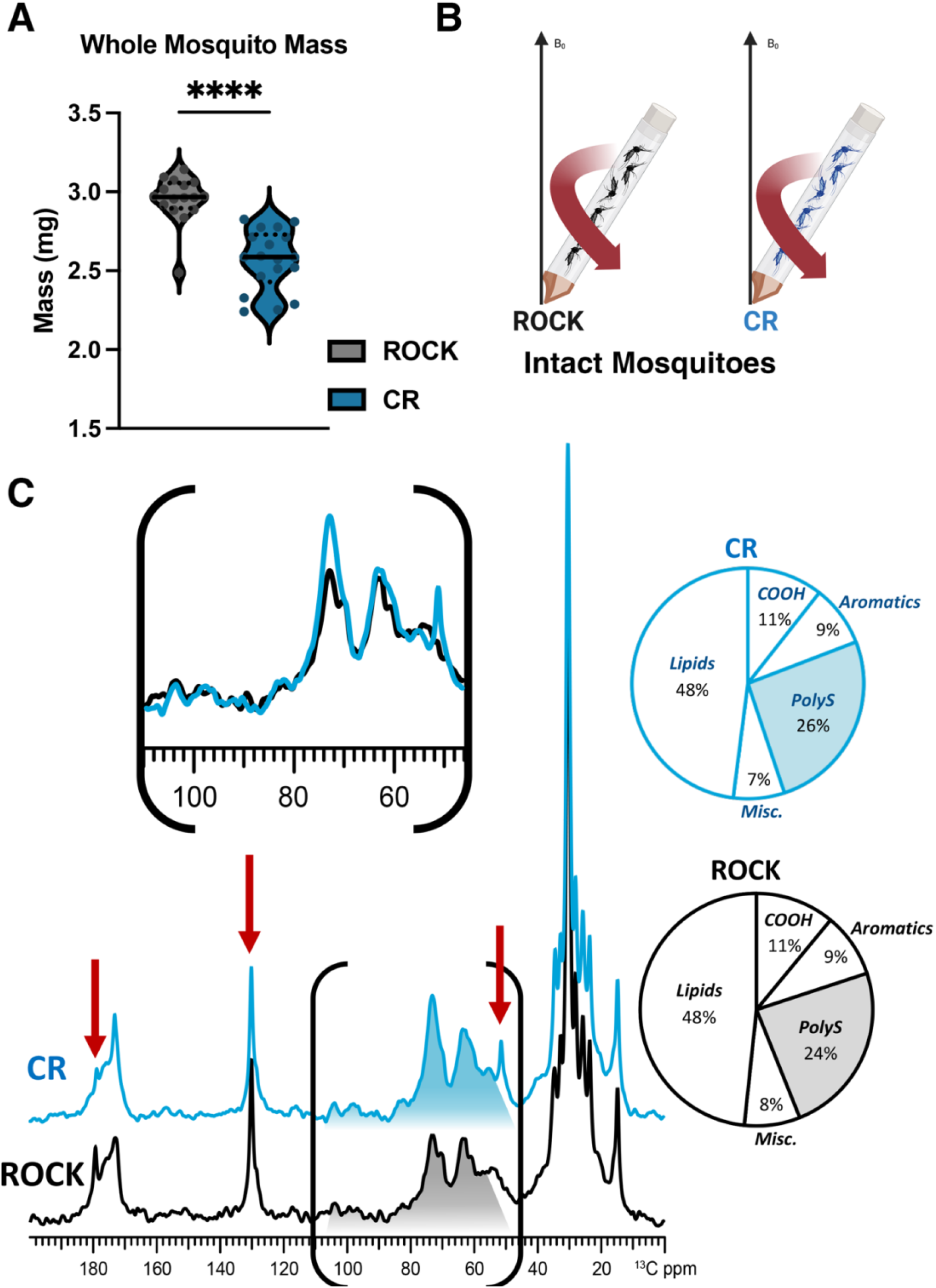
Solid-state NMR characterization of whole insecticide resistant and susceptible *Ae. aegypti*. **A:** Single mosquito mass estimated from pooled groups of 25 females across four biological replicates. Sample size: CR n = 16; ROCK n = 18. Mann-Whitney test p value = <0.0001 **B:** Schematic of material loaded into ssNMR rotor to compare female mosquitoes of both strains **C:** ^13^C DPMAS ssNMR (50-sec delay; quantitatively reliable) comparison for 20 whole female mosquitoes of each strain.

To explore this possibility, quantitatively reliable DPMAS ^13^C NMR experiments were performed on 20 whole CR or ROCK female mosquitoes (Fig. 2B). The spectra of CR (blue trace) and ROCK (black trace) whole mosquitoes are shown in Figure 2C, each normalized to the tallest peak (∼30 ppm). In contrast with the acid-resistant ssNMR data, the CR and ROCK whole-mosquito spectra contain more subtle differences in several spectral regions. Although it is visually apparent that the total NMR signal intensity displayed within the polysaccharide spectral region (∼55-110 ppm) is greater in the CR mosquito spectrum in comparison to the ROCK spectrum, quantitative analysis revealed that there is only a marginal difference between the two strains in terms of polysaccharide composition (26% vs 24% for CR and ROCK, respectively).

However, there are three notable differences between the CR and ROCK whole-mosquito spectra; namely, two sharp signals at ∼180, and 130 ppm that appear with greater intensity in the ROCK spectrum, and a sharp signal at 50 ppm that is displayed only in the CR spectrum. Specifically, the more prominent 180 ppm peak visible in the ROCK spectrum is where the carboxyl carbon of free fatty acids resonate, and a slightly more prominent peak at 130 ppm, which is typical for aromatic and ethylene carbons of unsaturated lipids [62]. In contrast, the identification of the peak at 50 ppm is more ambiguous: the chemical shift is consistent with a secondary or tertiary carbon that is covalently bonded to one or more electronegative atoms, such as oxygen or nitrogen. The intense signal at 50 ppm that appears only in the CYP spectrum could be present as a direct outcome of CYP gene overexpression. Since this CR strain is known to hydroxylate pyrethroid insecticides, in the absence of these compounds CYP enzymes might non-specifically hydroxylate structurally similar compounds; the aliphatic carbons of secondary metabolites that have been hydroxylated would resonate at 50 ppm.

### Changes in cuticular thickness over time

Cuticle thickening is an established cuticular resistance phenotype [18]. However, past work has not explored if cuticle thickness increases over time. Cross-sections of female CR and ROCK femurs were used to compare cuticle thickness and thickening over time at 3-5 d and 7-10 d post-eclosion from the same rearing cohort (Fig 3A, Sup. Fig. 2). Cross-sections were imaged using TEM. Total cuticle, endocuticle, and exocuticle measurements were obtained from 10 femurs per group (40 total femurs) and 10 images per femur were measured. CR resistant females had significantly larger total cuticle width than ROCK at both time points (Fig 3B, left). Notably, CR endocuticle width increased over time, whereas the ROCK endocuticle width did not increase. Moreover, cuticular thickening occurred primarily at the endocuticle (Fig 3B, middle). Analysis of TEM images showed that the exocuticle remained unchanged in both ROCK and CR when each strain was compared at 3-5 d and 7-10 d, thus serving as an internal control for consistent location of leg sectioning between samples (Fig 3B, right). Although exocuticle measurements of CR at 7-10 d (mean = 1.81 SD = 0.32) are statistically significantly larger than ROCK at 7-10 d (mean = 1.724 SD = 0.29), the 0.084 μm difference between mean exocuticle thicknesses is unlikely to be biologically significant.

**Figure 3:**
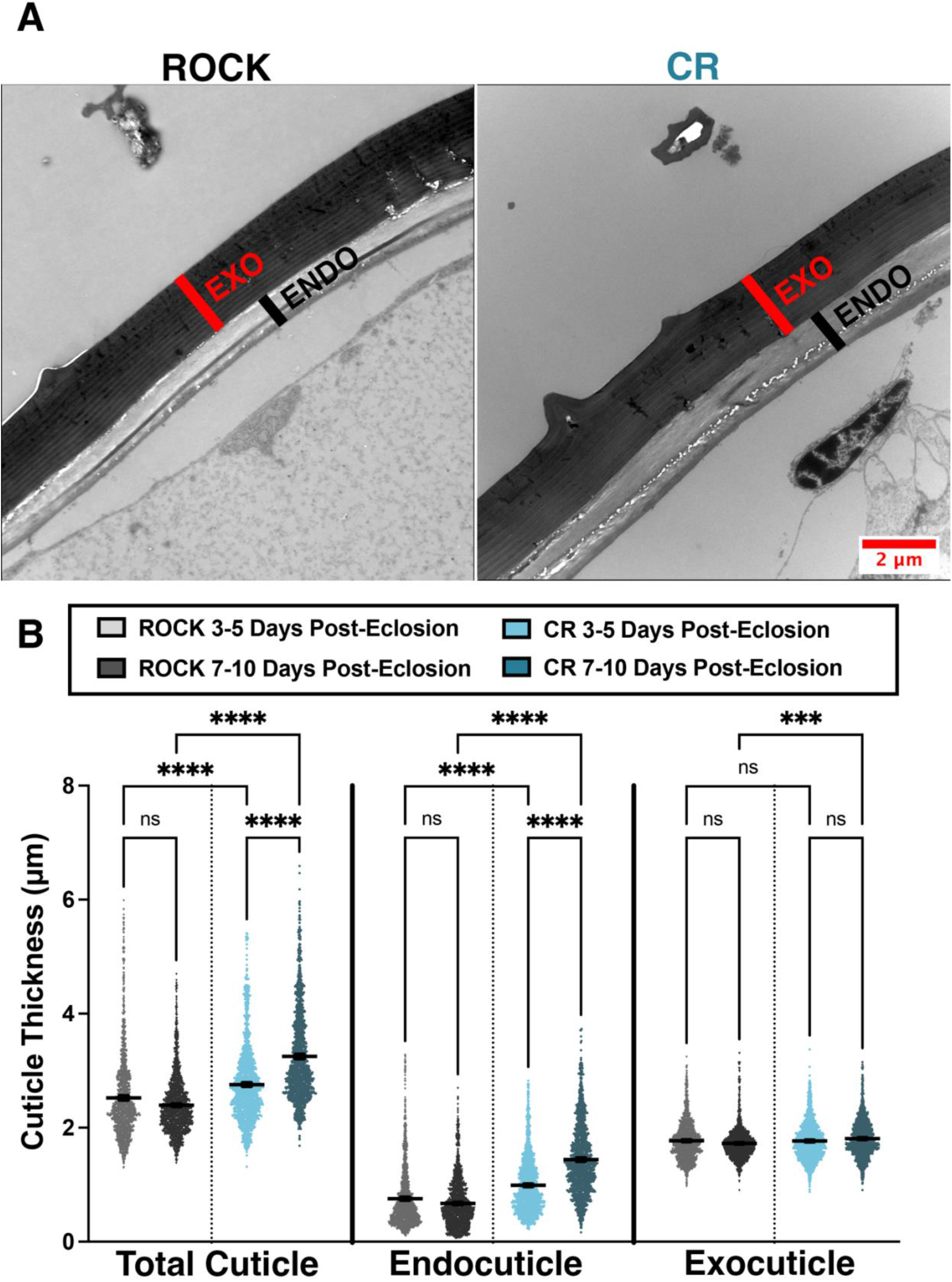
Time-course measurements of insecticide resistant and susceptible *Ae. aegypti* cuticular thickness. **A:** TEM images of 7–10 d ROCK (left) and CR (right) femurs indicating the exocuticle (red) and endocuticle (black). **B:** Cuticle measurements taken from cross sections of CR and ROCK female mosquito femurs both 3-5 d and 7-10 d post-eclosion. Data represent measurements from 10 femurs per strain per time point (40 total femurs) with 10 representative images analyzed per female. A minimum of 10 total cuticle, exocuticle, and endocuticle measurements were taken per image. Mean and 95% CI are shown on the graph. 3-5 d post-eclosion ROCK (light grey) N = 1,534 measurements; 3-5 d post-eclosion CR (light blue) N = 1,525 measurements. 7-10 d post-eclosion ROCK (dark grey) N = 1,688 measurements; 7-10 d post-eclosion CR (dark blue) N = 1,461 measurements. Kruskal-Wallis Test with Dunn’s multiple comparison. **** = <0.0001 *** = 0.0002.

### GC-MS analysis of cuticular hydrocarbons from CR and ROCK males and females

Increased abundance of cuticular lipids is correlated with insecticide resistance in the lab and field [46]. In the acid-resistant material of the insecticide resistant strain, we noted that the relative increase of the polysaccharide NMR spectral region coincided with a relative decrease in the lipid region. We were curious if this would correspond to a decrease in cuticular lipids. To investigate the abundance and composition of CR and ROCK cuticular hydrocarbon materials, we extracted the hydrocarbons from 3-5 d old CR and ROCK males and females. Hexane extractions were performed on groups of 20 pooled mosquitoes per extraction with an internal pentadecane standard. Samples were collected from three separate rearing cohorts. Total mosquitoes were as follows: ROCK females n = 180, ROCK males n = 140, CR females n = 140, CR males = 180. To estimate cuticular hydrocarbon (CHC) abundance, samples were referenced to the internal pentadecane standard. Total abundances were estimated from the sum of peak areas and divided by sample mass. GC-MS analysis of the extractions found that males and females of the ROCK and CR strains had similar alkane composition (Fig. 4A, 4B) and total abundance of CHCs (Fig. 4C).

**Figure 4:**
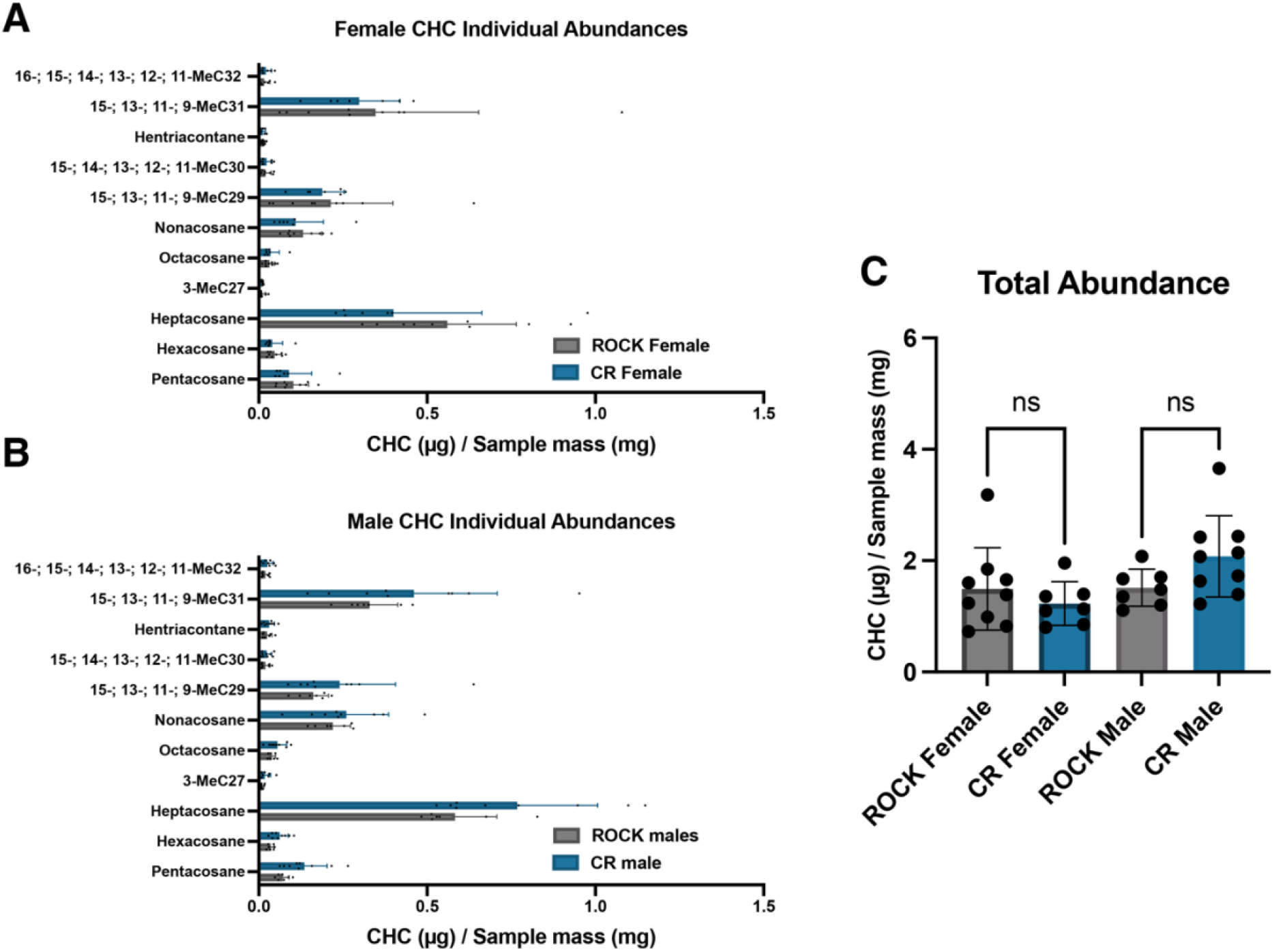
Cuticular hydrocarbon abundance and composition of resistant and susceptible *Ae. aegypti* males and females. **A:** Abundances of cuticular alkanes extracted from ROCK (grey) and CR (blue) females identified with GC-MS. Error bars represent mean with SD. **B:** Abundances of cuticular alkanes extracted from ROCK (grey) and CR (blue) males identified with GC-MS. Error bars represent mean with SD. **C:** Abundances of cuticular hydrocarbons extracted from ROCK (grey) and CR (blue) females and males estimated by summing peak area and adjusting by sample weight. All data points represent abundances from extractions of groups of 20 whole mosquitoes from 3 separate rearing cohorts.

**Figure 5:**
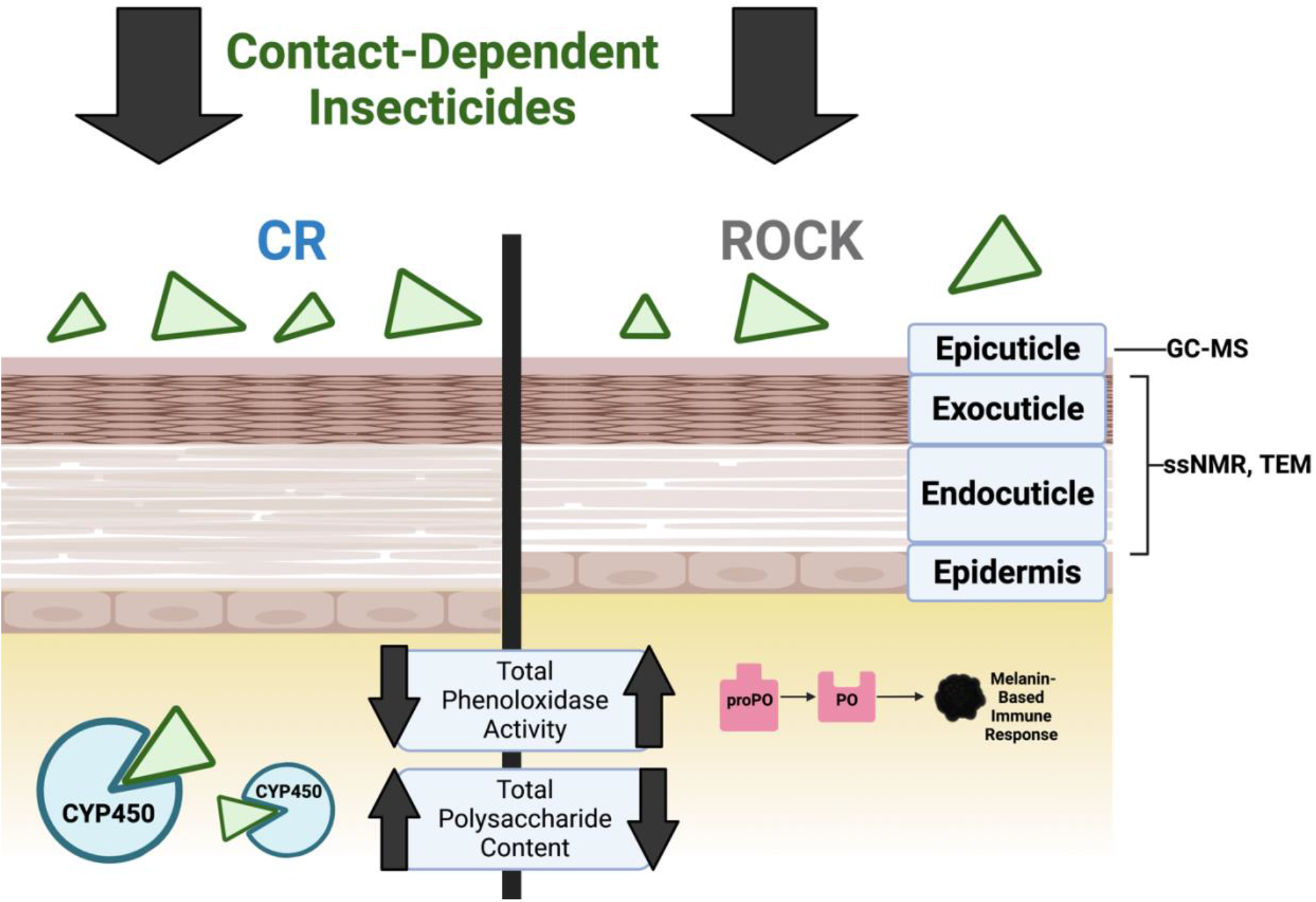
Schematic of cuticular structure and summary of methods used. Figure adapted from: Balabanidou et. al, 2018 [18]. Created with BioRender.com.

## Discussion

Cuticular insecticide resistance refers to insect cuticle alterations that reduce insecticide penetration, simultaneously limiting internal damage and increasing efficacy of internal physiological resistance mechanisms such as CYP enzyme detoxification and insecticide target-site mutations conferring KDR phenotypic resistance [18]. Compared to internal physiological mechanisms, cuticular resistance is much less well understood. Insect cuticle is essential for structural integrity, barrier protection, sensation, hydration, and chemical communication [19, 20]. Therefore, cuticular alterations may have unanticipated consequences in other aspects of insect physiology that require further study. Notably, past studies examining cuticular resistance have compared unrelated strains or strains with undefined resistance mechanisms. Comparing congenic strains that differ only in the trans-regulation of CYP gene overexpression supports the premise that our findings are related, either directly or indirectly, to the defined CYP-mediated resistance mechanisms and not purely a result of the genetic background. Previously, the SP strain, which possesses both a KDR resistance loci and the same metabolic resistance loci also present in the CR strain, was found to not be resistant to radiolabeled permethrin penetration after application to the notum [16]. However, this work did not characterize cuticle ultrastructure or composition [16]. Interactions between KDR and CYP resistance loci have been shown to impact physiology and influence fitness-costs [48, 49]. This is the case in the strain used in this study; isolation of the CR strain’s resistance loci from the KDR resistance loci resulted in greater fitness costs associated with CYP-resistance, including a reduced lifespan [49]. Therefore, it is possible that the cuticular changes we observed are related to the overexpression of CYP genes or associated with the isolation of the resistance loci in the CR strain. Future work comparing the cuticles of congenic strains with dual and isolated resistance loci would begin to answer these questions.

As noted previously, the cuticle is comprised of cuticular hydrocarbon-based waxes, chitin polysaccharides, cuticular proteins, and the phenolic biopolymers sclerotin and melanin. In contrast to other cuticular constituents, the contributions of sclerotin and melanin to cuticular resistance have been unexplored. This is primarily due to their relatively low abundance in the cuticle and amorphous structure. Within the cuticle, moreover, sclerotin and melanin are attached to other moieties via strong covalent bonds and are resistant to acid digestion [25, 57, 63]. In fungi, analysis of acid resistant material makes it possible to characterize not only fungal melanin, but components protected by melanin [55, 60, 64-66]. Adapting acid-digestion protocols used for fungal melanin enrichment allowed us to characterize the abundance and composition of associated material in the resistant cuticle [67]. While ssNMR comparison of ROCK and CR acid-resistant material showed no differences in melanin and sclerotin deposition correlating with CR resistance at baseline, the strains were not pre-exposed to insecticides. Therefore, the possibility that melanin or other phenolic compounds play a role in detoxification during direct exposure cannot be excluded.

Sclerotin and melanin play vital roles in several essential physiological processes including coloration, cuticular structural integrity, immunity, and wound healing [25-29]. Melanin produced by phenoloxidase is especially important in the insect immune response [26, 28, 29, 33]. Phenoloxidases exhibit broad substrate specificity and are able to metabolize a wide variety of phenolic-containing compounds, such as those secondary metabolites produced by plants, which are in turn toxic to insects if ingested in sufficient quantities [34, 35]. As such, these enzymes have a proposed role in detoxification in addition to immunity. Prior studies have demonstrated a potential link between increased phenoloxidase activity and insecticide resistant lab strains of the common house mosquito, *Culex pipiens*, although this trend was not apparent in resistant field populations [52, 53]. In contrast with the *C. pipiens* lab strains, our observation that the total phenoloxidase activity (estimated by *a*-chymotrypsin activation) was significantly decreased in the CR strain does not support the notion that phenoloxidases are playing a major role in detoxification. However, the biological relevance of this decrease during immune challenge is unknown.

Whereas ssNMR analysis of the acid-resistant material ruled out increased deposition of phenolic groups in the cuticle, it revealed higher relative polysaccharide content in the CR strain. The primary polysaccharide contributor could be identified as chitin, a premise supported because of acid resistance conferred by this material’s crosslinking ability to other cuticular moieties [30, 57-59, 61] and characteristic 55-ppm NMR resonance from the C2 ring carbon that is amide bonded to an acetyl group. This signal was clearly observed to display greater intensity in the CR sample spectrum (Fig. 1D inset). Nonetheless, it was unknown whether this increase in chitin corresponded to increased cuticle thickness, a commonly observed phenotype of cuticular resistance. By comparing the leg femur thickness of the CR insecticide-resistant *Ae. aegypti* strain to the congenic susceptible ROCK strain, we were able to demonstrate time-dependent cuticular thickening in the CR strain. CR females had significantly larger total cuticle and endocuticle thickness at both time points. Interestingly, the endocuticle thickness of CR females increased over time. Due to the timing of exocuticle formation, exocuticle thickness could serve as an internal control for consistent TEM sectioning location; the exocuticle thickness did not increase significantly over time in either strain.

Increased epicuticular hydrocarbon deposition is another documented cuticular resistance phenotype [44, 46, 47]. In contrast with the literature, GC-MS profiling of hexane-extracted hydrocarbons from 3–5-day-old males and females of both strains showed no significant differences in total cuticular hydrocarbon abundance or any identified individual alkanes. Nonetheless, we observed significant variability between samples, attributable to the three separate rearing cohorts, that rendered these comparisons challenging. To put these findings into context, it should be noted that several families of CYP genes are known to contribute to metabolic resistance. Populations of *An. arabiensis* and *An. gambiae* with overexpression of *CYP4G16* and *CYP4G17* exhibit increased cuticular hydrocarbon deposition associated with insecticide resistance and mating success [44, 46, 68]. Consistent with this literature, the insecticide resistant *Ae. aegypti* CR strain used in this study relies on overexpression of genes in the families CYP6 and CYP9, and not on CYP4G subfamily overexpression for conferral of synthetic pyrethroid resistance [69].

Although we observed no difference in cuticular hydrocarbon abundance, these lipids are known to play a role in mating of many insect species [46, 70, 71]. Previous work has demonstrated a mating defect associated with the resistance locus of the CR strain [49]. In a mating competition assay using congenic strains, susceptible ROCK males were more successful at mating with fellow susceptible ROCK females than males that exhibited both *kdr* and CYP resistance mechanisms [49]. Conversely, resistant females showed no preference between resistant and susceptible males [49]. Taken together, these results indicate that CYP resistance mechanisms may both reduce male mating fitness and alter the ability of resistant females to distinguish between potential mates [49]. In *Ae. aegypti*, the cuticle is an essential interface for chemosensory communication during mating and female choice is a primary driver of mating success [72, 73]. Recent work in *Ae. aegypti* has demonstrated that cuticular contact during mating induces chemosensory gene expression changes that are important in mate choice [73]. Due to the importance of cuticular contact in mating, the cuticular thickening we observed in the CR strain may reduce or alter chemosensory communication during mating, contributing to the mating defects associated with this metabolic resistance loci in lieu of hydrocarbon differences.

ssNMR has been performed previously on whole mosquitoes to show global metabolic changes [62]. As the CR strain is metabolically insecticide resistant, we aimed to gain more detailed insights into the cellular architecture and overall molecular composition of the whole mosquito by probing for global spectral changes. When we performed ssNMR analysis on intact female mosquitoes, the CR strain exhibited a marginally greater polysaccharide content in comparison to ROCK. We observed the same trend when comparing the polysaccharide content of acid-resistant material yielded by the two strains. In contrast to acid-resistant material, whole organisms contain a wide variety of polysaccharides that exhibit only minor differences in chemical shift, which results in significant spectral overlap and thus precludes the identification of specific types of polysaccharides. Therefore, it cannot be ruled out that the marginally greater polysaccharide content of CYP whole mosquitoes is reflective of the increased chitin content observed in the acid-resistant material.

Alternatively, it is possible that CR mosquitoes exhibit an increase in polysaccharide content as a direct result of CYP gene overexpression. In insects, a well-established mechanism of detoxification is the conjugation of xenobiotics to glucose by glucosyl transferases [68]. The proposed mechanism of CYP resistance in this strain is the hydroxylation of aromatic ring carbons of pyrethroids; the hydroxylated compounds are further conjugated with glucosides or amino acids to mediate excretion of these polar secondary metabolites [16, 74]. Thus, this detoxification mechanism requires the availability of readily mobilizable carbohydrate reserves, which could potentially contribute to the increased polysaccharide content observed in this strain.

In addition to the overall increase of relative NMR signal intensity in the polysaccharide region, the CR spectrum displayed a prominent sharp peak at ∼50 ppm that was not observed in the ROCK spectrum. CYPs participate in the detoxification of xenobiotics such as chemical insecticides (e.g., pyrethroids) via the hydroxylation of key sites, which increases their polarity and in turn promotes their excretion. Metabolically resistant mosquitoes exhibit constitutive overexpression of CYPs; thus, when reared in a lab setting in the absence of any xenobiotics, increased CYP activity could result in the hydroxylation of structurally similar non-toxic moieties. Although not fully understood, CYPs and other downstream detoxifying enzymes are thought to preferentially hydroxylate the arene and aliphatic carbons of pyrethroids [75]. Depending on the local chemical environment, certain hydroxylated aliphatic carbons resonate at ∼50 ppm and thus could plausibly explain the unique appearance of this peak in the CR whole mosquito spectrum.

Further comparison of the CR and ROCK whole-mosquito data revealed additional spectral differences which could potentially reflect a difference in metabolic state between the two strains. Namely, there are two sharp signals at ∼180 and 130 ppm that are of greater intensity in the ROCK spectrum. The peaks at 180 and 130 ppm are characteristic of rapidly-tumbling unsaturated free fatty acids found in fat bodies [76-78]: the signal at 180 ppm is unambiguously attributable to the carbon within a carboxylic acid group of an unesterified fatty acid [76, 79], whereas the signal at 130 ppm is attributable to the olefinic carbons of fatty acids containing at least one point of unsaturation [76, 79], which are predominant in fat bodies [80] and preferentially liberated from triacylglycerols as compared with saturated fatty acids [81]. Taken together, our data suggest that ROCK mosquitoes have a greater content of free fatty acids in comparison to the CYP strain. Notably, these differences in free fatty acid peak intensities were not observed in the spectra of the acid-resistant material, which suggests they reflect a change in global metabolism rather than a change in cuticular molecular architecture. Free fatty acids are an important energy source in insect metabolism [69]; they are stored in fat bodies in the form of triacylglycerols and are liberated immediately prior to utilization for energy production. Thus, the comparatively lower content of fatty acids observed in CR mosquitoes offers evidence of the high energetic cost of maintaining CYP resistance [82, 83]. The high energy demands of the CYP strain and overexpression of detoxification mechanisms are likely to require mobilization of carbohydrate reserves, resulting in chitin production and therefore causing cuticle thickening to correlate with metabolic resistance. We hypothesize that global metabolic changes impacting carbohydrate metabolism contribute indirectly to the cuticular alterations often observed in resistant strains, rather than a distinct selected phenomenon. Indeed, our observation of endocuticle thickening over time supports this notion.

Our work profiling the contributions of polysaccharide, lipid, and phenolic biopolymers to cuticular resistance revealed cuticular changes in the CR strain. While TEM allowed us to monitor the cuticle ultrastructure for differences, a combination of solid-state Nuclear Magnetic Resonance (ssNMR) and Gas Chromatography Mass Spectrometry (GC-MS) yielded insights into cuticular composition. We were additionally able to rule out baseline differences in both cuticular hydrocarbons and phenolic biopolymer deposition between CR and ROCK. However, we observed endocuticular thickening over time and an associated increased polysaccharide content in both acid-resistant cuticular material and whole CR mosquito. Nevertheless, the ability of cuticle alterations to act synergistically with other resistance mechanisms and impact important processes such as mating emphasizes the importance of better understanding the biomolecules that contribute to cuticular insecticide resistance in *Ae. aegypti* and related vectors that jeopardize human health.

## Acknowledgements

We thank Barbara Smith of the Johns Hopkins Microscope Core Facility for expert assistance with TEM imaging and sample sectioning. Thank you to Scott Lab members Juan Silva and Cera Fisher for sending the CR strain. We are grateful to Doug Norris for manuscript advice and encouragement. We appreciate the help of Quigly Dragotakes and Daniel Smith in manuscript editing.

## Funding Acknowledgements

This work was supported by the National Institutes of Health, grant number R01-AIxxxyyy (E.J., OTHERS, A.C.). E.J was also supported by the National Institutes of Health, grant number T32-AI138953-03. C.C. was also supported by the Brescia Fund of the Department of Chemistry and Biochemistry at The City College of New York.

## Author contributions

E.J: conceptualization, methodology, validation, formal analysis, investigation, writing-original draft, writing-review and editing, visualization. C.C: methodology, validation, formal analysis, investigation, writing-original draft, writing-review and editing, visualization. S.R.T: methodology, validation, formal analysis, investigation, writing-review and editing, visualization. M.W: supervision, project administration, writing-review, and editing. E.C: methodology, writing-review, and editing. J.G.S: Resources, supervision, writing-review, and editing. N.A.B: Resources, supervision, writing-review, and editing. C.J.M: Resources, supervision, validation, writing-review, and editing. R.E.S: Resources, supervision, writing-review and editing, and funding acquisition. A.C: conceptualization, supervision, project administration, resources, supervision, writing-review and editing, and funding acquisition.

## Methods

### Mosquito strains

The congenic *Ae. aegypti* CR and ROCK strains used in this study were produced and supplied by Dr. Jeff Scott’s laboratory at Cornell University (Ithaca, New York). CYP resistance alleles originating from the resistant Singapore (SP) strain were introgressed into the ROCK background to produce a strain with only CYP-mediated resistance that was congenic to the well characterized insecticide susceptible Rockefeller (ROCK) strain [17, 49, 84, 85].

### Mosquito rearing

The *Ae. aegypti* CR and ROCK strains were maintained with a 12 h light:dark photoperiod at 27°C and 80% relative humidity. Larvae were fed 1 pellet of Cichlid Gold® Fish Food (Hikari, Himeji, Japan) per 50 larvae [69]. Eggs were vacuum hatched in a 1L flask in diH_2_O for 30 minutes to maximize synchronized development. After 24 h, larvae were sorted to a density of 200 larvae/1 L of diH_2_O. After eclosion, mosquitoes were maintained on 10% sucrose using cotton wicks.

### *Drosophila melanogaster* strains and rearing

The *D. melanogaster* strains used in this study, WT-CantonS (BDSC 64349), ebonyS (BDSC 498), and *yw* (BDSC 1495) were obtained from the Bloomington Drosophila Stock Center (Bloomington, IN). All *D. melanogaster* strains were maintained on our standard fly food [86] on a 12 h light:dark photoperiod at 25°C and 80% relative humidity. Adult flies (4-7 day old) were collected and sexed for further processing. Acid-resistant material was collected from groups of 25 homogenized female flies. Samples were homogenized in 500 μL diH_2_O using an electric pellet pestle cordless motor (Kimble). After homogenization, 500 μL of 12M HCl was added to the homogenate (6M final concentration). Samples were digested for 24 hours on an Eppendorf ThermoMixer C shaker at 85°C with 700 RPM shaking. After digestion, samples were spun down for 30 minutes at room temperature at 15,000 RCF. Samples were washed three times, first with 1 mL 1X PBS, then 1 mL 10% PBS, and finally 1 mL diH_2_O. Washed melanin samples were lyophilized, weighed, and combined for ssNMR analysis.

### Phenoloxidase activity

To measure phenoloxidase activity, single mosquitoes were crushed in 35 μL of cold PBS using an electric pellet pestle cordless motor (Kimble) with ScienceWare Disposable Polypropylene Pestles (VWR Catalogue # 66001-104). After a 2-minute 3,000 RPM spin at 4°C, 15 μL of hemolymph homogenate was recovered and frozen on dry ice for 2 minutes for hemocyte lysis. Samples were stored at -80°C until analysis. Hemolymph homogenate samples were thawed on ice. In a transparent, flat bottom 96-well plate 5 μL of hemolymph homogenate was mixed with 20 μL PBS, 20 μL 20 mM L-DOPA (4mg/mL) (3,4-Dihydroxy-L-phenylalanine, Sigma Catalogue # D9628-25G), and 140 μL diH_2_O with or without 0.07 mg/mL *a*-chymotrypsin (Worthington Biochemical, Catalog #LS001432). Melanization activity was determined through SpectraMax iD5 spectrophotometer readings at 492 nm 30°C for 45 minutes with 1 reading/minute [53].

### Acid digestion and mosquito weights

Female mosquitoes were collected 5-7 d post-eclosion and weighed in groups of 25 in 1.5 mL microcentrifuge tubes. The weight of single mosquitoes was estimated from these measurements to reduce error. To obtain acid resistant material, 25 female mosquitoes were homogenized in 500 μL diH_2_O using an electric pellet pestle cordless motor (Kimble). After homogenization, 500 μL of 12M HCl was added to the homogenate (6M final concentration). Samples were digested for 24 hours on an Eppendorf ThermoMixer C shaker at 85°C with 700 RPM shaking. After digestion, samples were spun down for 30 minutes at room temperature at 15,000 RCF. Samples were washed three times, first with 1 mL 1X PBS, then 1 mL 10% PBS, and finally 1 mL diH_2_O. Washed melanin samples were lyophilized, weighed, and combined for ssNMR analysis.

### Solid-state NMR spectroscopy

Solid-state NMR experiments were conducted on a Varian (Agilent) DirectDrive2 (DD2) spectrometer operating at a 1H frequency of 600 MHz and equipped with a 1.6-mm T3 HXY fastMAS probe (Agilent Technologies, Santa Clara, CA); all measurements were carried out using a magic-angle spinning (MAS) rate of 15.00 ± 0.02 kHz at a spectrometer-set temperature of 25 °C. Data were obtained on ∼X-Y mg of lyophilized sample mass yielded by HCl hydrolysis of each CR and ROCK female mosquitoes to analyze the acid-resistant material. To analyze the whole mosquitos, data were obtained on 20 intact lyophilized female mosquitoes from either the CR or ROCK strain, equivalent to approximately XYZ mg. Both sets of samples were examined using ^13^C direct-polarization (DPMAS) experiments conducted with 90° pulse lengths of 1.2 and 1.4 μs for 1H and ^13^C, respectively; 104-kHz heteronuclear decoupling using the small phase incremental alternation pulse sequence (SPINAL) was applied during signal acquisition. The DPMAS experiments used a long recycle delay (50-s) to generate spectra with quantitatively reliable signal intensities. Thus, the relative amounts of carbon-containing constituents present in the samples could be estimated using the GNU image manipulation program (GIMP) by measuring the integrated signal intensity within the spectral region corresponding to each moiety and comparing it to the total integrated signal intensity of the spectrum.

### TEM sectioning and image analysis

Groups of 10 midlegs were removed from ROCK and CR females of the same rearing cohort at 3-5 d post-eclosion and 7-10 d post-eclosion (40 total midlegs). Midlegs were severed at the tibia just after the femur with a razor blade and were fixed in 2.5% glutaraldehyde, 3 mM MgCl2, and 0.1 M sodium cacodylate (pH 7.2) overnight at 4 °C.

After buffer rinse, samples were postfixed in 1% osmium tetroxide, 1.25% potassium ferrocyanide in 0.1 M sodium cacodylate for at least 1 h (no more than two) on ice in the dark. After the fixing step, samples were rinsed in dH2O, followed by uranyl acetate (2%, aq.) (0.22 μm filtered, 2.5 h, dark), dehydrated in a graded series of ethanol and embedded in Spurrs (Electron Microscopy Sciences) resin. Samples were polymerized at 60 °C overnight. Thin sections, 60 to 90 nm, were cut with a diamond knife on a Leica UCT ultramicrotome and picked up with 2 × 1 mm Formvar copper slot grids. To obtain consistent comparable segments across samples, femur cross-sections were obtained 200 nm into the femur from the direction of the tibia. Grids were stained with 2% aqueous uranyl acetate followed by lead citrate and observed with a Hitachi 7600 TEM at 80 kV. Images were captured at 10,000X magnification using an AMT CCD XR80 (8-megapixel side mount AMT XR80 high-resolution, high-speed camera).

Ten representative images were chosen per femur. Each image captured ∼18 μm of total cuticle length. ≤10 measurements were taken per image using ImageJ2 version 2.30/1.53f. Measurements of total cuticle and endocuticle were taken at the same point and exocuticle was calculated by subtracting these values. Areas of the cuticle containing structural modifications were excluded from measurements.

### Cuticular hydrocarbon extraction

Pools of 20 mosquitoes were collected at 3-5 d post-eclosion, weighed, and stored in glass vials at -20°C until sample processing. Samples were collected from three separate rearing cohorts. The pools of 20 mosquitoes were submerged for 30 min in 400 μL GC-MS quality hexane and 16 µg/sample pentadecane internal standard. Each extraction was purified through a column chromatography quality Silica gel (Pore Size 60 Å 0.063-0.200 mm) to final a volume of 1.5 mL of hexane and evaporated with N_2_ gas. After evaporation, samples were frozen at -20°C until further analysis.

### GC-MS Analysis

Samples were analyzed by gas chromatography/mass spectrometry (7890B GC, 5977N MSD, Agilent, USA). Concentrated hydrocarbons were resuspended in 30 μL of hexane and 1 μL of each sample was injected onto a HP-5MS capillary column (30 m length x 25 mm diameter x 0.25 µm film thickness). The GC oven was programmed with an initial temperature of 50 °C with a 2 min hold followed by an increase of 20 °C/min to 300 °C with a 6 min hold. A helium carrier gas with a flow rate of 1.2 mL/min^-1^ was used. The MS analyzer was set to acquire over a range of m/z 35-500 and was operated in EI mode. The ion source and transfer line were set to 230 °C and 300 °C respectively. Analyte peak areas were normalized to the internal standard and by sample weights. Compound identification was achieved by comparison of mass spectra with the NIST Mass Spectral Library version 2.2 and retention time matching with analytical reference standards.

### Statistical analysis

Data were analyzed with Prism Version 9.3.1. Figure 1A left: The data were normally distributed (Shapiro-Wilk test) and a two-tailed unpaired t-test was performed. Figure 1A right: The data were not normally distributed (Shapiro-Wilk test) and a two-tailed Mann-Whitney test was performed. Figure 1B: The data were normally distributed (Shapiro-Wilk test) and a two-tailed unpaired t-test was performed. Figure 2A: The data were not normally distributed (Shapiro-Wilk test) and a two-tailed Mann-Whitney test was performed. Figure 3B: The data were not normally distributed (Kolmogorov–Smirnov test) and a Kruskal-Wallis test with Dunn’s multiple comparisons test was performed with 12 comparisons.

---

**Supplementary Figure 1.**
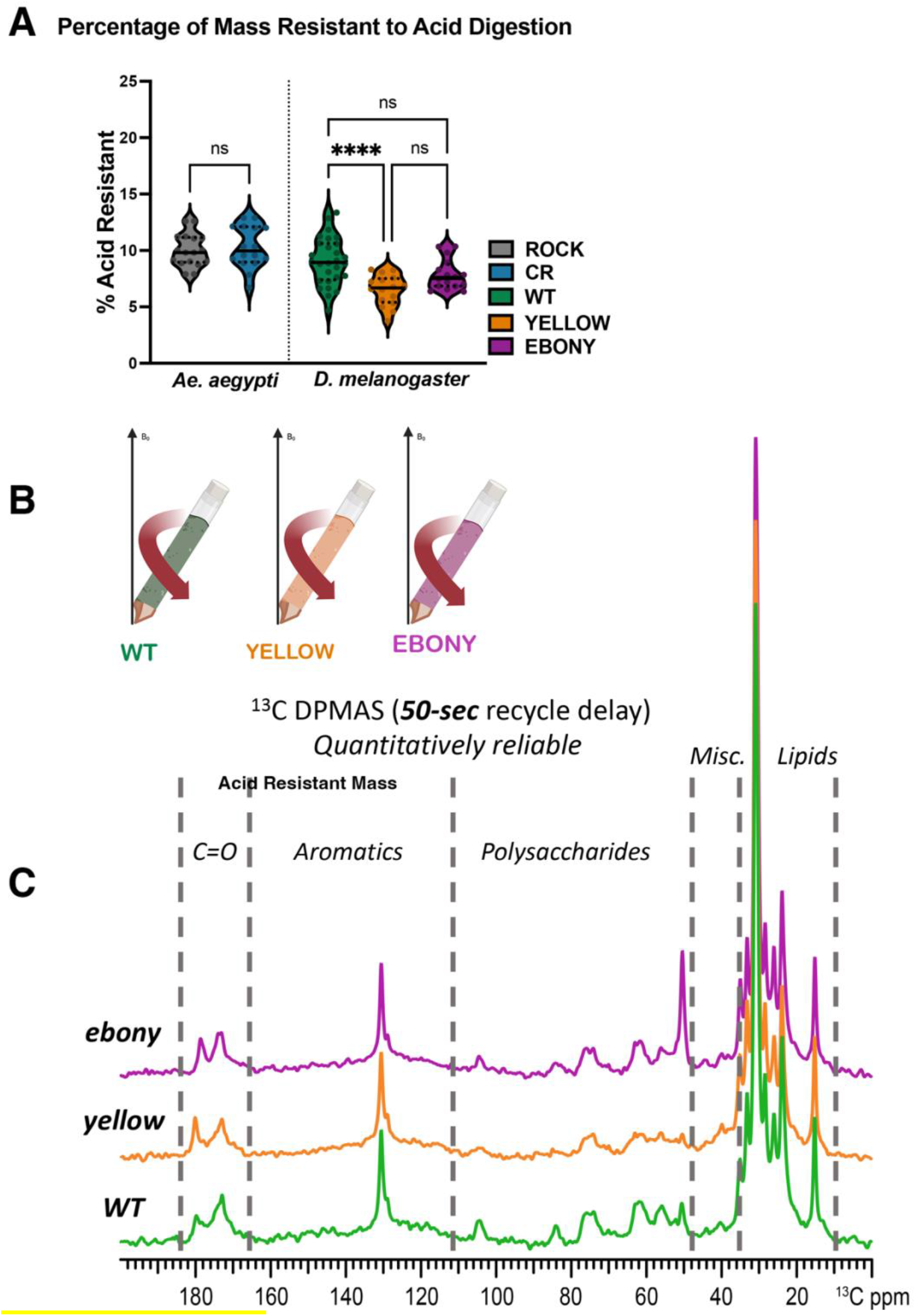
**A:** Percentage of female *Ae. aegypti* ROCK (grey) and CR (dark blue) and *D. melanogaster* WT (green), yellow (orange), and ebony (pink) wet weights that were resistant to acid digestion. All digestion samples contained 25 females each across three pooled biological replicates. Sample number: CR n = 16, ROCK n = 17, WT n= 28, yellow n= 16, Ebony n = 16. One-way ANOVA with Tukey’s Multiple Comparison test p value: ****= <0.0001, ** = 0.0048 **B:** Schematic of material loaded into ssNMR rotor to compare acid-resistant material from *D. melanogaster* strains **C:** direct-polarization (DPMAS) Carbon-13 (^13^C) ssNMR (50-sec delay; quantitatively reliable) comparison of acid-resistant material of the WT (green), yellow (orange), and ebony (pink) strains pooled from three biological replicates.

**Supplementary Figure 2.**
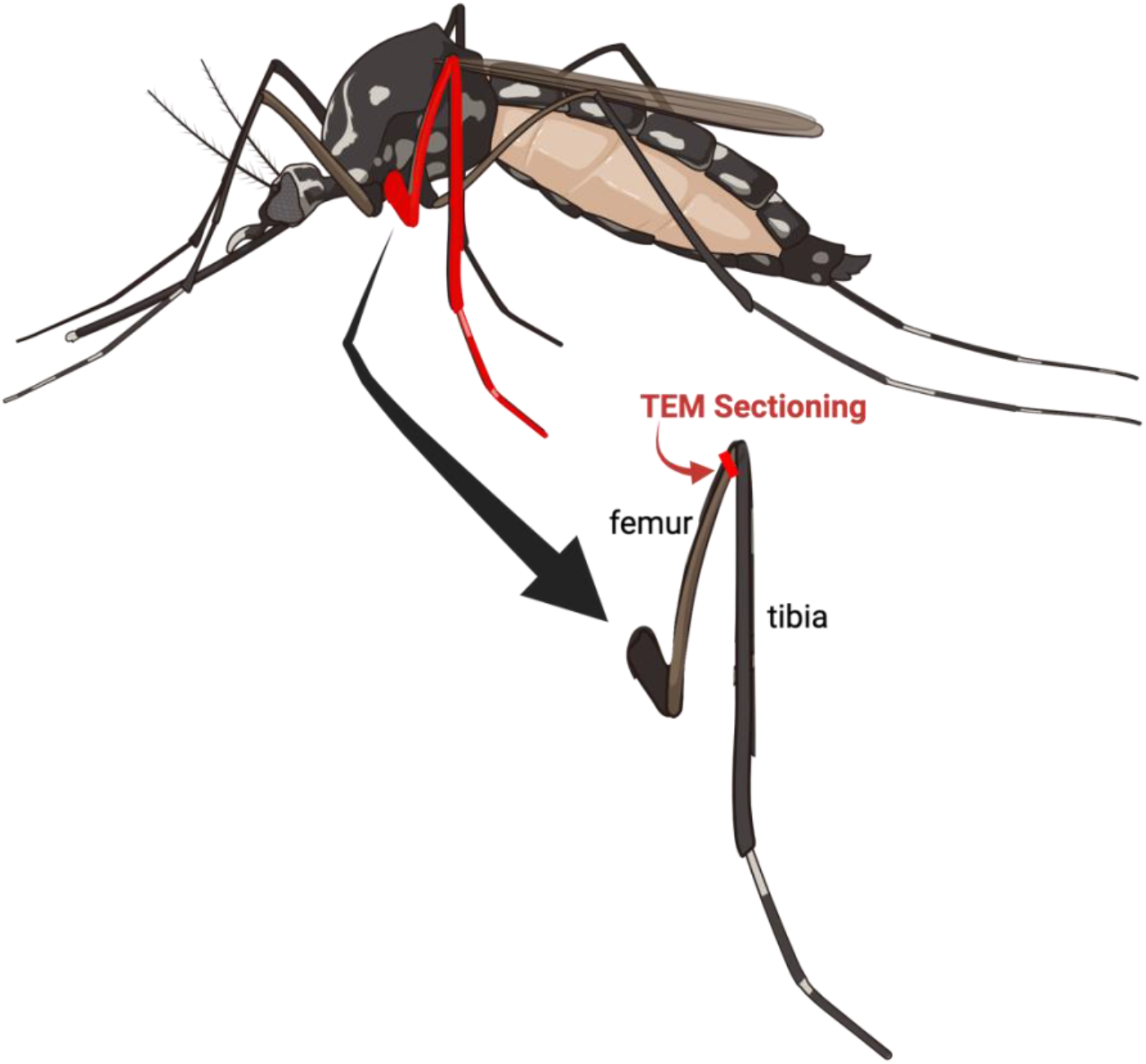
**A:** Schematic of TEM sectioning performed 200 nm into the midleg femur.

## References

1. Vega-Rúa, A., et al., Chikungunya virus transmission potential by local Aedes mosquitoes in the Americas and Europe. PLoS Negl Trop Dis, 2015. 9(5): p. e0003780.

2. Shepard, D.S., et al., The global economic burden of dengue: a systematic analysis. Lancet Infect Dis, 2016. 16(8): p. 935–41.

3. Bhatt, S., et al., The global distribution and burden of dengue. Nature, 2013. 496(7446): p. 504–7.

4. Messina, J.P., et al., The current and future global distribution and population at risk of dengue. Nature Microbiology, 2019. 4(9): p. 1508–1515.

5. Marcombe, S., et al., Pyrethroid resistance reduces the efficacy of space sprays for dengue control on the island of Martinique (Caribbean). PLoS Negl Trop Dis, 2011. 5(6): p. e1202.

6. Maciel-de-Freitas, R., et al., Undesirable consequences of insecticide resistance following Aedes aegypti control activities due to a dengue outbreak. PLoS One, 2014. 9(3): p. e92424.

7. Moyes, C.L., et al., Contemporary status of insecticide resistance in the major Aedes vectors of arboviruses infecting humans. PLoS Negl Trop Dis, 2017. 11(7): p. e0005625.

8. Rahman, R.U., et al., Insecticide resistance and underlying targets-site and metabolic mechanisms in Aedes aegypti and Aedes albopictus from Lahore, Pakistan. Scientific Reports, 2021. 11(1): p. 4555.

9. Zalucki, M.P. and M.J. Furlong, Behavior as a mechanism of insecticide resistance: evaluation of the evidence. Current Opinion in Insect Science, 2017. 21: p. 19–25.

10. Sparks, T.C., et al., The role of behavior in insecticide resistance. Pesticide Science, 1989. 26(4): p. 383–399.

11. Scott, J.G. Investigating Mechanisms of Insecticide Resistance: Methods, Strategies, and Pitfalls. 1990.

12. Ingham, V.A., et al., A sensory appendage protein protects malaria vectors from pyrethroids. Nature, 2020. 577(7790): p. 376–380.

13. Ensley, S.M., Chapter 39 - Pyrethrins and Pyrethroids, in Veterinary Toxicology (Third Edition), R.C. Gupta, Editor. 2018, Academic Press. p. 515–520.

14. Dong, K., et al., Molecular biology of insect sodium channels and pyrethroid resistance. Insect Biochem Mol Biol, 2014. 50: p. 1–17.

15. Hemingway, J., et al., The molecular basis of insecticide resistance in mosquitoes. Insect Biochem Mol Biol, 2004. 34(7): p. 653–65.

16. Kasai, S., et al., Mechanisms of pyrethroid resistance in the dengue mosquito vector, Aedes aegypti: target site insensitivity, penetration, and metabolism. PLoS Negl Trop Dis, 2014. 8(6): p. e2948.

17. Smith, L.B., et al., CYP-mediated resistance and cross-resistance to pyrethroids and organophosphates in Aedes aegypti in the presence and absence of kdr. Pestic Biochem Physiol, 2019. 160: p. 119–126.

18. Balabanidou, V., L. Grigoraki, and J. Vontas, Insect cuticle: a critical determinant of insecticide resistance. Current Opinion in Insect Science, 2018. 27: p. 68–74.

19. Wigglesworth, V.B., THE INSECT CUTICLE. Biological Reviews, 1948. 23(4): p. 408–451.

20. Vincent, J.F.V., Arthropod cuticle: a natural composite shell system. Composites Part A: Applied Science and Manufacturing, 2002. 33(10): p. 1311–1315.

21. Fine, B.C., P.J. Godin, and E.M. Thain, Penetration of Pyrethrin I labelled with Carbon-14 into Susceptible and Pyrethroid Resistant Houseflies. Nature, 1963. 199(4896): p. 927–928.

22. Forgash, A.J., B.J. Cook, and R.C. Riley, Mechanisms of Resistance in Diazinon-Selected Multi-Resistant Musca domestica1. Journal of Economic Entomology, 1962. 55(4): p. 544–551.

23. Andersen, S.O., Biochemistry of Insect Cuticle. Annual Review of Entomology, 1979. 24(1): p. 29–59.

24. Muthukrishnan, S., et al., Insect Cuticular Chitin Contributes to Form and Function. Current pharmaceutical design, 2020. 26(29): p. 3530–3545.

25. Sugumaran, M., Molecular mechanisms for mammalian melanogenesis. Comparison with insect cuticular sclerotization. FEBS Lett, 1991. 295(1-3): p. 233–9.

26. Christensen, B.M., et al., Melanization immune responses in mosquito vectors. Trends in Parasitology, 2005. 21(4): p. 192–199.

27. Whitten, M.M.A. and C.J. Coates, Re-evaluation of insect melanogenesis research: Views from the dark side. Pigment Cell & Melanoma Research, 2017. 30(4): p. 386–401.

28. Nappi, A.J. and B.M. Christensen, Melanogenesis and associated cytotoxic reactions: applications to insect innate immunity. Insect Biochem Mol Biol, 2005. 35(5): p. 443–59.

29. Sugumaran, M., Comparative biochemistry of eumelanogenesis and the protective roles of phenoloxidase and melanin in insects. Pigment Cell Res, 2002. 15(1): p. 2–9.

30. Duplais, C., et al., Gut bacteria are essential for normal cuticle development in herbivorous turtle ants. Nature Communications, 2021. 12(1): p. 676.

31. Andersen, S.O., Insect cuticular sclerotization: A review. Insect Biochemistry and Molecular Biology, 2010. 40(3): p. 166–178.

32. González-Santoyo, I. and A. Córdoba-Aguilar, Phenoloxidase: a key component of the insect immune system. Entomologia Experimentalis et Applicata, 2012. 142(1): p. 1–16.

33. Söderhäll, K. and L. Cerenius, Role of the prophenoloxidase-activating system in invertebrate immunity. Current Opinion in Immunology, 1998. 10(1): p. 23–28.

34. Wu, K., et al., Plant phenolics are detoxified by prophenoloxidase in the insect gut. Scientific Reports, 2015. 5(1): p. 16823.

35. Liu, S., et al., Does phenoloxidase contributed to the resistance? Selection with butane-fipronil enhanced its activities from diamondback moths. Open Biochem J, 2009. 3: p. 9–13.

36. Luna-Acosta, A., et al., Enhanced immunological and detoxification responses in Pacific oysters, Crassostrea gigas, exposed to chemically dispersed oil. Water Research, 2011. 45(14): p. 4103–4118.

37. Yu, J.-J., et al., Effects of piperonyl butoxide synergism and cuticular thickening on the contact irritancy response of field Aedes aegypti (Diptera: Culicidae) to deltamethrin. Pest Management Science, 2021. 77(12): p. 5557–5565.

38. Samal, R.R. and S. Kumar, Cuticular thickening associated with insecticide resistance in dengue vector, Aedes aegypti L. International Journal of Tropical Insect Science, 2021. 41(1): p. 809–820.

39. Noppun, V., T. Saito, and T. Miyata, Cuticular penetration of S-fenvalerate in fenvalerate-resistant and susceptible strains of the diamondback moth, Plutella xylostella (L.). Pesticide Biochemistry and Physiology, 1989. 33(1): p. 83–87.

40. Balabanidou, V., et al., Mosquitoes cloak their legs to resist insecticides. Proceedings of the Royal Society B: Biological Sciences, 2019. 286(1907): p. 20191091.

41. Wood, O., et al., Cuticle thickening associated with pyrethroid resistance in the major malaria vector Anopheles funestus. Parasit Vectors, 2010. 3: p. 67.

42. Yahouédo, G.A., et al., Contributions of cuticle permeability and enzyme detoxification to pyrethroid resistance in the major malaria vector Anopheles gambiae. Scientific Reports, 2017. 7(1): p. 11091.

43. Bass, C. and C.M. Jones, Mosquitoes boost body armor to resist insecticide attack. Proceedings of the National Academy of Sciences, 2016. 113(33): p. 9145–9147.

44. Balabanidou, V., et al., Cytochrome P450 associated with insecticide resistance catalyzes cuticular hydrocarbon production in Anopheles gambiae. Proc Natl Acad Sci U S A, 2016. 113(33): p. 9268–73.

45. Li, D.-T., et al., Ten fatty acyl-CoA reductase family genes were essential for the survival of the destructive rice pest, Nilaparvata lugens. Pest Management Science, 2020. 76(7): p. 2304–2315.

46. Adams, K.L., et al., Cuticular hydrocarbons are associated with mating success and insecticide resistance in malaria vectors. Communications Biology, 2021. 4(1): p. 911.

47. Qiu, Y., et al., An insect-specific P450 oxidative decarbonylase for cuticular hydrocarbon biosynthesis. Proc Natl Acad Sci U S A, 2012. 109(37): p. 14858–63.

48. Hardstone, M.C., C.A. Leichter, and J.G. Scott, Multiplicative interaction between the two major mechanisms of permethrin resistance, kdr and cytochrome P450-monooxygenase detoxification, in mosquitoes. J Evol Biol, 2009. 22(2): p. 416–23.

49. Smith, L.B., et al., Fitness costs of individual and combined pyrethroid resistance mechanisms, kdr and CYP-mediated detoxification, in Aedes aegypti. PLOS Neglected Tropical Diseases, 2021. 15(3): p. e0009271.

50. Smith, L.B., et al., CYP-mediated permethrin resistance in Aedes aegypti and evidence for trans-regulation. PLOS Neglected Tropical Diseases, 2018. 12(11): p. e0006933.

51. Wang, Y., P. Aisen, and A. Casadevall, Melanin, melanin “ghosts,” and melanin composition in Cryptococcus neoformans. Infect Immun, 1996. 64(7): p. 2420–4.

52. Vézilier, J., et al., The impact of insecticide resistance on Culex pipiens immunity. Evolutionary Applications, 2013. 6(3): p. 497–509.

53. Cornet, S., S. Gandon, and A. Rivero, Patterns of phenoloxidase activity in insecticide resistant and susceptible mosquitoes differ between laboratory-selected and wild-caught individuals. Parasites & Vectors, 2013. 6(1): p. 315.

54. Cerenius, L., B.L. Lee, and K. Söderhäll, The proPO-system: pros and cons for its role in invertebrate immunity. Trends in Immunology, 2008. 29(6): p. 263–271.

55. Chrissian, C., et al., Solid-state NMR spectroscopy identifies three classes of lipids in Cryptococcus neoformans melanized cell walls and whole fungal cells. J Biol Chem, 2020. 295(44): p. 15083–15096.

56. Latocha, M., et al., Pyrolytic GC-MS analysis of melanin from black, gray and yellow strains of Drosophila melanogaster. Journal of Analytical and Applied Pyrolysis, 2000. 56(1): p. 89–98.

57. Christensen, A.M., et al., Detection of cross-links in insect cuticle by REDOR NMR spectroscopy. Journal of the American Chemical Society, 1991. 113(18): p. 6799–6802.

58. Chrissian, C., et al., Unconventional Constituents and Shared Molecular Architecture of the Melanized Cell Wall of C. neoformans and Spore Wall of S. cerevisiae. J Fungi (Basel), 2020. 6(4).

59. Chatterjee, S., et al., Using solid-state NMR to monitor the molecular consequences of Cryptococcus neoformans melanization with different catecholamine precursors. Biochemistry, 2012. 51(31): p. 6080–8.

60. Chrissian, C., et al., Melanin deposition in two Cryptococcus species depends on cell-wall composition and flexibility. J Biol Chem, 2020. 295(7): p. 1815–1828.

61. Kramer, K.J., T.L. Hopkins, and J. Schaefer, Applications of solids NMR to the analysis of insect sclerotized structures. Insect Biochemistry and Molecular Biology, 1995. 25(10): p. 1067–1080.

62. Chang, J., et al., Solid-state NMR reveals differential carbohydrate utilization in diapausing Culex pipiens. Scientific Reports, 2016. 6(1): p. 37350.

63. Sugumaran, M. and H. Barek, Critical Analysis of the Melanogenic Pathway in Insects and Higher Animals. International Journal of Molecular Sciences, 2016. 17(10): p. 1753–n/a.

64. Chatterjee, S., et al., Solid-state NMR Reveals the Carbon-based Molecular Architecture of Cryptococcus neoformans Fungal Eumelanins in the Cell Wall. J Biol Chem, 2015. 290(22): p. 13779–90.

65. Camacho, E., et al., The structural unit of melanin in the cell wall of the fungal pathogen Cryptococcus neoformans. J Biol Chem, 2019. 294(27): p. 10471–10489.

66. Baker, R.P., et al., Cryptococcus neoformans melanization incorporates multiple catecholamines to produce polytypic melanin. J Biol Chem, 2022. 298(1): p. 101519.

67. Zhong, J., et al., Following fungal melanin biosynthesis with solid-state NMR: biopolymer molecular structures and possible connections to cell-wall polysaccharides. Biochemistry, 2008. 47(16): p. 4701–10.

68. Jones, C.M., et al., The dynamics of pyrethroid resistance in Anopheles arabiensis from Zanzibar and an assessment of the underlying genetic basis. Parasites & Vectors, 2013. 6(1): p. 343.

69. Sun, H., et al., Transcriptomic and proteomic analysis of pyrethroid resistance in the CKR strain of Aedes aegypti. PLOS Neglected Tropical Diseases, 2021. 15(11): p. e0009871.

70. Savarit, F., et al., Genetic elimination of known pheromones reveals the fundamental chemical bases of mating and isolation in Drosophila. Proc Natl Acad Sci U S A, 1999. 96(16): p. 9015–20.

71. Polerstock, A.R., S.D. Eigenbrode, and M.J. Klowden, Mating Alters the Cuticular Hydrocarbons of Female Anopheles gambiae sensu stricto and Aedes aegypti (Diptera: Culicidae). Journal of Medical Entomology, 2002. 39(3): p. 545–552.

72. Aldersley, A. and L.J. Cator, Female resistance and harmonic convergence influence male mating success in Aedes aegypti. Scientific Reports, 2019. 9(1): p. 2145.

73. Alonso, D.P., et al., Gene expression profile of Aedes aegypti females in courtship and mating. Scientific Reports, 2019. 9(1): p. 15492.

74. Shono, T., T. Unai, and J.E. Casida, Metabolism of permethrin isomers in American cockroach adults, house fly adults, and cabbage looper larvae. Pesticide Biochemistry and Physiology, 1978. 9(1): p. 96–106.

75. Stevenson, B.J., et al., Pinpointing P450s associated with pyrethroid metabolism in the dengue vector, Aedes aegypti: developing new tools to combat insecticide resistance. PLoS Negl Trop Dis, 2012. 6(3): p. e1595.

76. Alexandri, E., et al., High Resolution NMR Spectroscopy as a Structural and Analytical Tool for Unsaturated Lipids in Solution. Molecules, 2017. 22(10).

77. Hakumäki, J.M. and R.A. Kauppinen, 1H NMR visible lipids in the life and death of cells. Trends Biochem Sci, 2000. 25(8): p. 357–62.

78. Rémy, C., et al., Evidence that mobile lipids detected in rat brain glioma by 1H nuclear magnetic resonance correspond to lipid droplets. Cancer Res, 1997. 57(3): p. 407–14.

79. Rakhmatullin, I.Z., et al., NMR chemical shifts of carbon atoms and characteristic shift ranges in the oil sample. Petroleum Research, 2022. 7(2): p. 269–274.

80. Stadler Martin, J., Lipid composition of fat body and its contribution to the maturing oöcytes in Pyrrhocoris apterus. Journal of Insect Physiology, 1969. 15(6): p. 1025–1045.

81. Arrese, E.L., et al., Lipid storage and mobilization in insects: current status and future directions. Insect Biochem Mol Biol, 2001. 31(1): p. 7–17.

82. Rivero, A., et al., Energetic cost of insecticide resistance in Culex pipiens mosquitoes. J Med Entomol, 2011. 48(3): p. 694–700.

83. Hardstone, M.C., et al., Differences in development, glycogen, and lipid content associated with cytochrome P450-mediated permethrin resistance in Culex pipiens quinquefasciatus (Diptera: Culicidae). J Med Entomol, 2010. 47(2): p. 188–98.

84. Smith, L.B., S. Kasai, and J.G. Scott, Pyrethroid resistance in Aedes aegypti and Aedes albopictus: Important mosquito vectors of human diseases. Pesticide Biochemistry and Physiology, 2016. 133: p. 1–12.

85. Kuno, G., Early history of laboratory breeding of Aedes aegypti (Diptera: Culicidae) focusing on the origins and use of selected strains. J Med Entomol, 2010. 47(6): p. 957–71.

86. Lesperance, D.N.A. and N.A. Broderick, Meta-analysis of Diets Used in Drosophila Microbiome Research and Introduction of the Drosophila Dietary Composition Calculator (DDCC). G3 Genes|Genomes|Genetics, 2020. 10(7): p. 2207–2211.

